# ALIGN: A modular framework for quantitative analysis of longitudinal development in *Caenorhabditis elegans*

**DOI:** 10.64898/2026.11.03.763986

**Authors:** Sacha Psalmon, Benjamin D. Towbin

**Affiliations:** Institute of Cell Biology, University of Bern, Bern, Switzerland; Graduate School for Cellular and Biomedical Sciences, University of Bern, Bern, Switzerland

**Keywords:** growth, development, *C. elegans*, live imaging, automatic image analysis

## Abstract

Longitudinal live imaging is a powerful tool for studying growth, development and behaviour. However, reproducible and scalable analysis of the large resulting datasets remains a major bottleneck. Here, we present ALIGN (Automated Longitudinal Image-based Growth aNalyzer), a modular framework that integrates deep learning-based segmentation, automated quality control, computational straightening, feature extraction and interactive annotation within a single configurable pipeline that can be operated end-to-end by users without coding expertise. Applying ALIGN to longitudinal development of *C. elegans*, we demonstrate accurate segmentation across heterogeneous imaging conditions, detection of larval stage transitions from growth trajectories, and data-driven quality control, ensuring high measurement precision. Computational straightening and alignment enable consistent analysis of motile animals, removing motion artefacts from three-dimensional optical sectioning. Applying ALIGN to measure pharyngeal pumping and growth rates in wild-type and eat-2 mutants uncovered a non-proportional relationship between feeding and growth. ALIGN thus provides generalizable tools and resources for longitudinal, single-animal analysis of development.

## Introduction

Affordable and programmable microscopy has made time-lapse imaging broadly accessibl e across the life sciences (Diederich et al., 2020; Li et al., 2020), enabling longitudinal observation of biological systems in vivo at high throughput and individual resolution. These advances in data acquisition are further leveraged by recent progress in computational analysis, including machine and deep learning methodologies (Moen et al., 2019), which greatly improve the prediction of future trajectories from past observations, opening up a wide range of applications from fundamental research in biology to medical diagnostics (Buggenthin et al., 2017; Whitwell et al., 2020).

The nematode *C. elegans* is particularly well suited for longitudinal studies due to its small size, optical transparency, and short lifecycle. These features enable parallel monitoring of hundreds of individual animals from embryogenesis to adulthood over days to weeks (Turek et al., 2015). A wide range of experimental platforms, including microfluidic devices (Keil et al., 2017; Berger et al., 2021) and sealed cultivation chambers (Turek et al., 2015), support continuous imaging of individual animals. However, a major remaining bottleneck in turning the large-scale datasets generated by these approaches into biological insight is their reproducible and reliable processing, their quantitative interpretation, and the management of their storage.

Specifically, several key challenges continue to limit the widespread adoption of longitudinal imaging workflows: First, data processing pipelines are often fragmented and lack standardisation, limiting reproducibility across experiments and laboratories. Second, while deep learning-based image analysis has been widely adopted, trained models frequently fail to generalise across imaging conditions, requiring substantial retraining even after minor changes in acquisition settings. Third, integrating segmentation, quality control, feature extraction, and downstream analysis into a unified workflow remains technically demanding, particularly for non-specialists. Even as large language model (LLM)-based code generation lowers the technical barrier to writing analysis scripts, the need for standardised, reproducible, and auditable workflows remains an urgent and timely problem.

Here, we address these challenges by developing ALIGN (Automated Longitudinal Image-based Growth aNalyzer), a modular and reproducible workflow framework for the analysis of long-term live imaging data with application to *C. elegans*. ALIGN integrates a complete set of longitudinal image analysis steps, including segmentation, automated quality control, spatial alignment and straightening, feature extraction, and interactive annotation within a single configurable pipeline. By combining standardised data handling, scalable processing, and user-friendly interfaces, ALIGN enables reproducible analysis of large-scale imaging datasets while substantially reducing the need for programming expertise. This framework facilitates the application of advanced image analysis methods to longitudinal biological data and lowers the barrier for their adoption across diverse experimental settings.

## Results

### A modular framework for image processing and analysis of large time-lapse microscopy datasets

To create a reproducible and integrated workflow for longitudinal time-lapse microscopy, we developed ALIGN (Automated Longitudinal Image-based Growth aNalyzer), an open-source image analysis framework implemented in Python 3. ALIGN unifies data processing, model training, and interactive analysis within a single modular architecture. This design enables extension by expert users, while maintaining accessibility for users without extensive programming experience.

ALIGN is organised into four complementary components: (1) a Python package providing core methods for image processing and feature extraction; (2) a dedicated pipelining engine for executing complete analysis workflows; (3) a graphical user interface (GUI), implemented as a Shiny (The Shiny development team, 2026) application, for interactive visualisation, annotation, and correction of time series data; and (4) a collection of napari (napari contributors, 2019) plugins for efficient generation and curation of training datasets for machine learning-based analysis. Separation of core methods from workflow execution enables extensibility while supporting fully automated and reproducible analyses across datasets and laboratories. Integration of the GUI connects quantitative outputs directly to raw image data, enabling efficient validation and correction, while the napari plugins facilitate adaptation of prediction models by non-expert users. Unlike general-purpose workflow managers (Di Tommaso et al., 2017; Mölder et al ., 2025), ALIGN is specifically designed to integrate image analysis, machine learning, and interactive curation within a single reproducible framework.

Reproducibility in ALIGN is achieved by representing analysis workfl ows as explicit sequences of atomic tasks (called “blocks”). For example, a typical workflow may consist of sequential blocks for segmentation, straightening, quality control, feature extraction, and automatic event detection, followed by manual curation, and data plotting (Fig. 1). Workflows are defined through a human-readable and easily editable configuration file (written in YAML) with standardised input and output formats, ensuring consistent data flow between processing steps. Centralised parameter specification i s designed to yield identical results upon re-analysis and repeatable configurations across experiments. Because the configuration file constitutes the primary user interface, complete workflows can be constructed and executed without writing code.

**Figure 1:**
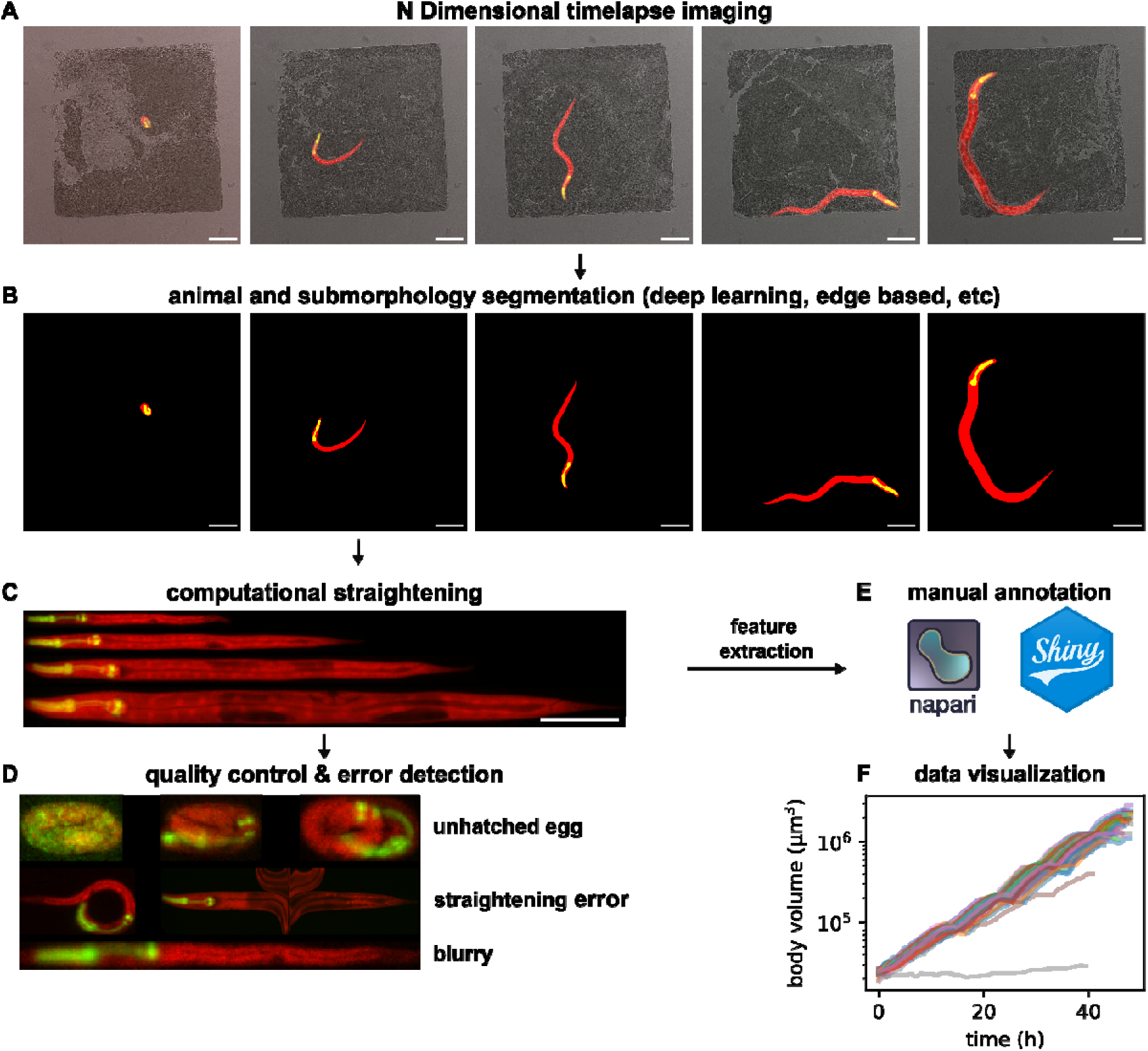
Overview of the ALIGN pipeline modules. **(A)** Example raw input data: Images of a worm trapped in a growth chamber at different stages of development from embryo to adulthood. Animals expressing GFP in the pharynx and mScarlet in all somatic tissues. (**B)** Deep learning-based segmentation of the images shown in (A). **(C)** Computational straightening and alignment of the images in (A), based on the segmentations shown in (B). **(D)** Examples of images flagged by automated quality control as unhatched embryos, poor image quality, or processing errors. **(E)** Interactive interfaces enable manual inspection and annotation. **(F)** Standardised output format ensures seamless and reproducible data visualisation through existing software packages. Shown is a time series of the body volume of 58 wild-type worms over the first 48h of their development after hatching. Scale bars: 100 µm.

Unlike existing general-purpose, graph-based workflow systems (Mölder et al., 2025), ALIGN adopts a linear and declarative execution model that avoids complex task dependency resolution while maintaining full transparency of processing steps. This design improves robustness and reduces computational overhead of the pipeline engine. To support large-scale datasets, ALIGN allows for automated task submission via SLURM for distributed resource management. Using modest resources (32 CPU cores, 1 consumer-grade GPU), analysis of full developmental time series of hundreds of individuals, summing up to over 100,000 images, can be completed overnight. Importantly, automated processing is tightly integrated with visualisation and manual curation, allowing users to inspect, validate, and refine resul ts within a unified environment.

In the following sections, we describe the individual components of ALIGN in the context of a representative workflow for analysing *C. elegans* development. This workflow includes segmentation of individual animals from time series data, computational straightening and alignment, automated and trainable quality control, three-dimensional reconstruction, and extraction of biologically relevant features such as larval stage transitions and organismal size. We further present tools for interactive annotation and correction, as well as workflows for training and adapting machine learning models for segmentation and quality control.

### Custom and pre-trained models enable robust and adaptable object segmentation

Accurate and reliable segmentation of organismal and cellular features across heterogeneous imaging conditions remains a major challenge in longitudinal microscopy, as models trained on one dataset often fail to generalise to new experimental settings. To address this, we implemented a flexible workflow for rapid generation and adaptation of segmentation models within ALIGN (Fig. 2A, 2B).

**Figure 2:**
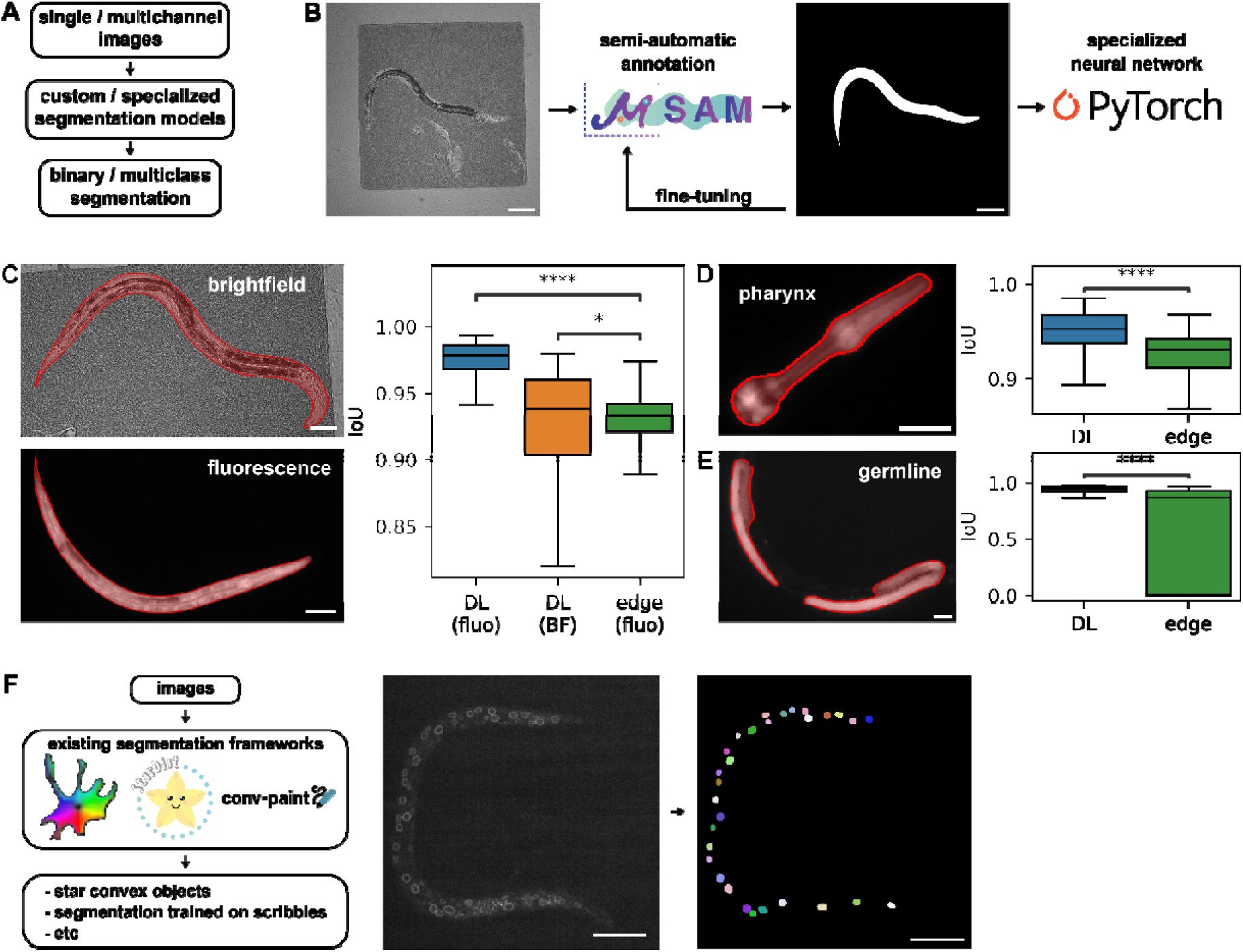
Comparison of segmentation algorithms implemented in ALIGN. **(A)** Modular segmentation workflow allows multimodal input and output. **(B)** The neural network training workflow uses user prompt-dependent microSAM segmentation to create training data for prompt-independent segmentation models. **(C)** Examples of whole worm body segmentation using deep learning (DL) segmentation models trained as shown in (B) from brightfield images (top) or from individuals ubiquitously expressing the fluorescent protein mScarlet (bottom, *eft-3*::mScarlet). Boxplot shows segmentation accuracy (IoU compared to masks created with the help of microSAM) of the two different methods: DL compared against edge-based (edge) segmentation of fluorescent signal (brightfield (BF)=295 images of 274 unique individuals sampled from 16 independent experiments, fluorescence (fluo)=201 images of 193 unique individuals from 30 independent experiments; DL (fluo) vs edge (fluo): Wilcoxon signed-rank test, two-sided (same images); DL (BF) vs edge (fluo): Mann-Whitney-Wilcoxon test, two-sided (disjoint image sets); no multiple-comparison corrections were applied). **(D)** Example of pharyngeal segmentation using a GFP marker expressed in the pharyngeal muscle (*myo-2p*::GFP). Boxplot of segmentation accuracy of trained model compared to edge-based segmentation (n=276 images of 257 individuals from 22 independent experiments; Wilcoxon signed-rank test, two-sided). **(E)** As (D), but for fluorescently labelled germline (*glh-1*p::GFP) (n=192 images of 172 unique individuals from 9 independent experiments; Wilcoxon signed-rank test, two-sided). **(F)** Existing segmentation workflows can be incorporated in the ALIGN pipeline. As an example, integration of StarDist for segmentation of nuclei is shown. Raw image (left) shows an individual expressing the nuclear envelope protein EMR-1 tagged with mCherry. Right image shows segmentation output by StarDist integrated in the ALIGN pipeline. Scale bars: 50 µm. *P<0.05; ****P<0.0001. Exact p-values provided in Supplemental Table 2. Boxplots: centre line, median; box, interquartile range (IQR); whiskers, 1.5×IQR.

ALIGN allows for the creation of balanced training and validation sets from a large pool of imbalanced data. A YAML configuration file lets users specify desired proportions of different classes of images, such as the developmental stage inspected, or the microscope used for the acquisition. ALIGN then automatically selects representative subsets of images that capture the diversity of experimental conditions or specimen features as specified by the user. These images can be annotated using microSAM (Archit et al., 2025; Kirillov et al., 2023), a prompt-based segmentation model that facilitates efficient generation of training labels (Fig. 2B). For complex tasks, the microSAM model can be iteratively refined after initial annotation of a few images to further reduce manual effort for creating training data (Fig. 2B). The resulting annotated dataset is used in ALIGN to train a U-Net-based segmentation model (Ronneberger et al., 2015). These models operate independently of prompt input and can be directly integrated into automated workflows (Fig. 2C-E). In summary, ALIGN uses microSAM to create ground truth data with minimal human prompting input. This data is then used to train a dedicated standalone neural network for prompt-free segmentation. An example of automated segmentation of arrayed individuals of *C. elegans* throughout development is shown in Supplemental Movie 1.

Using this workflow, high segmentation accuracy can be achieved even with limited training data. For the datasets presented here, fewer than 3,000 annotated individuals were sufficient to train U-Net models for segmentation of animals and fluorescently labelled substructures. Using microSAM (Archit et al., 2025; Kirillov et al., 2023) and the automated image selection by ALI GN, training data of this size can be generated within 6-12 hours by a single researcher. Upon need, the resulting base model can be fine-tuned with additional training data, e.g. i f segmentation performance drops due to sample treatments or illumination settings that substantially alter the appearance of the sample. Furthermore, when combining images from multiple microscopy systems, the trained models generalised to data acquired with different hardware configurations, including both high-end epifluorescence systems and lower-cost imaging setups (Li et al., 2020), with comparable performance (Fig. S1). In all tested cases, deep learning-based segmentation outperformed classical image analysis approaches using thresholding or edge segmentation (Table 1, Fig. 2C-E).

**Table 1:** Performance of ALIGN’s segmentation methods. Average IoU score of each method across a representative test dataset (body (fluorescence): n=201; body (brightfield): n=295; pharynx: n=276; germline: n=192).

|  | Whole body (fluorescence) | Whole body (brightfield) | Pharynx (fluorescence) | Germline (fluorescence) |
|---|---|---|---|---|
| U-Net++ (efficientnet-b5) | **0.975** | **0.911** | **0.948** | **0.926** |
| U-Net++ (efficientnet-b2) | 0.966 | 0.899 | 0.944 | 0.921 |
| Edge-based | 0.924 | 0.002 | 0.864 | 0.599 |

For conditions with weak contrast, such as segmentation of animals in dense bacterial backgrounds from transmitted light images, performance can be improved by increasing the size and diversity of the training dataset, reaching good levels of accuracy (0.975 vs 0.911 mean IoU score, Table 1, Fig. 2C). For near-perfect segmentation, fluorescence-based high-contrast images remain the preferred choice when technically feasible.

While we have found the above-described workflow to be effective for most applications, ALIGN also supports the integration of alternative pre-trained segmentation frameworks such as Cellpose (Pachitariu et al., 2025; Stringer et al., 2021) and StarDist (Schmidt et al ., 2018; Weigert et al., 2020) (Fig. 2F). Independent of the chosen model, ALIGN standardises pre- and post-processing, manages memory usage, and enables batch execution, thereby reducing the technical complexity associated with deploying deep learning-based segmentation methods. Together, ALIGN hence allows for rapid development and deployment of segmentation models that can be adapted to diverse experimental conditions without requiring expertise in neural network design or high-performance computing.

### Computational straightening enables quantitative analysis of motile animals

A prominent challenge of longitudinal imaging is that individual animals change in position, posture, and orientation between time points. This variability limits direct comparison of images across time and complicates extraction of morphological and dynamic features. In addition, movement during acquisition of three-dimensional image stacks leads to misalignment of optical sections, impairing accurate 3D reconstruction. For *C. elegans*, microfluidic immobilisation approaches have been used to mitigate these issues (Berger et al., 2021; Keil et al., 2017). However, physical immobilisation is labour-intensive, technically demanding, and difficult to scale. Moreover, applying mechanical force on worms may perturb their physiology.

To avoid these limitations, ALIGN implements a computational straightening and alignment procedure that reconstructs a consistent coordinate system for each animal across time points and imaging planes (Fig. 3). Building on previously described approaches (Peng et al., 2008), we optimised an algorithm that extracts the animal midline from the segmentation mask via Medial Axis Transform (MAT), followed by branch pruning and extension to the object boundaries (Fig. 3A). A coordinate system orthogonal to the midline is then computed and used to warp the image into a straightened representation (Fig. 3A).

**Figure 3:**
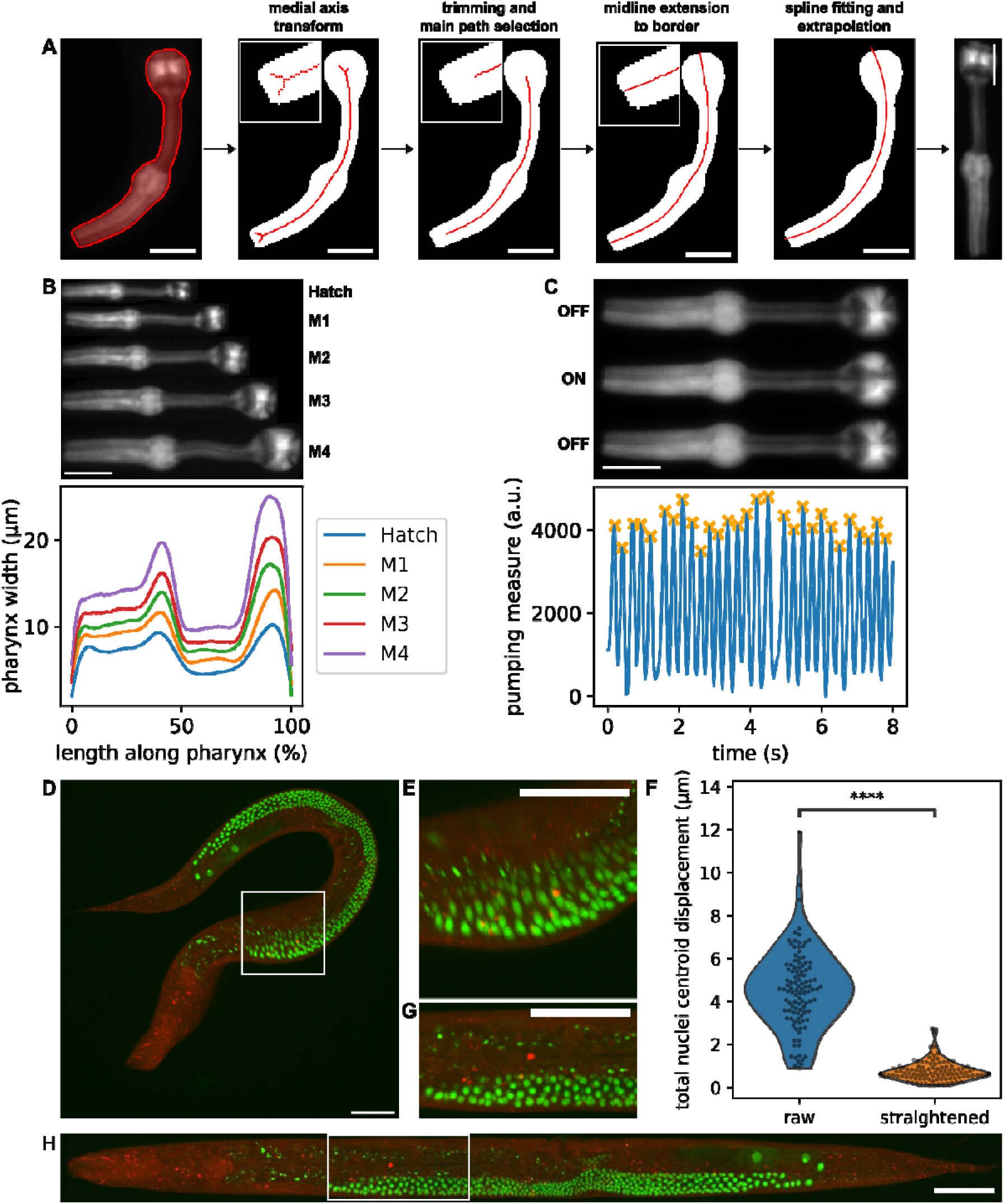
In-depth explanation of ALIGN’s computational image straightening. **(A)** Illustration of the straightening algorithm for a pharynx image and mask. The method finds the midline through MAT, trims branches and selects the main path, extends the midline to the border and fits the midline with a spline. **(B)** Straightened pharynxes of a worm at different larval stages and corresponding extracted width profiles. **(C)** Pharynx at different steps of food ingestion (pumping) and extracted pumping index used to compute feeding rate. **(D)** Maximum intensity projection of a confocal z-stack of a moving *C. elegans*. The strain expresses a ubiquitously expressed red fluorescent protein (*eft-3p*::mScarlet) and a germline-specific nuclear GFP marker (*Y75B12B.1p::nls*::GFP). **(E)** Magnified view of the area in (D) highlighted by the white rectangle where most of the movement occurs. **(F)** Violin plot of the XY displacement across the Z stack of the centroid of each individual nucleus represented in (E, raw) and (G, straightened). Nuclei were segmented using Cellpose and nuclear identity was propagated across the z-axis based on an overlap threshold across planes (threshold = 0.5 IoU). Nuclei that appeared for fewer than 5 planes were ignored from the rest of the analysis. Displacement is defined as the Euclidean distance between the centroid of the first occurrence of each nucleus and the last. (n=108 nuclei; Wilcoxon signed-rank test, two-sided). **(G)** Magnified view of the same part as (E) but taken from the reconstructed stack shown in (H). **(H)** Maximum intensity projection of the same stack as in (D), but 3D reconstructed after straightening and alignment. Scale bars: 25 µm (A-C) and 50 µm (D-H). ****P<0.0001. Exact p-values provided in Supplemental Table 2.

This transformation enables direct and consistent quantification of morphological features, such as length and width profiles, across time and individuals (Fig. 3B, Supplemental Movie 2). In addition, straightened representations facilitate visualisation and analysis of dynamic developmental processes. As one application, we implemented automated extraction of pharyngeal pumping rates from high-frame-rate recordings acquired longitudinally throughout development (Fig. 3C), building on methods previously applied to adult animals (Bonnard et al., 2022).

The straightening approach further extends to three-dimensional datasets, where animal movement between optical sections leads to misalignment of z-stacks (Fig. 3D-E). To address this, adjacent planes are iteratively aligned using phase cross-correlation prior to final warping, enabling reconstruction of coherent three-dimensional representations of motile animals (Fig. 3G-H). As a proof of concept, we applied this approach to the 3D reconstruction of GFP-labelled germ cell nuclei from a moving animal. Whereas the unaligned image displayed a systematic drift in the positioning of the centre of mass of the same nucleus imaged at different planes due to animal movement, this drift was reduced more than 6-fold after reconstruction (Fig. 3F).

Beyond enabling quantitative analysis, straightening also restricts analysis to the region of the image containing the object of study, rather than the entire field of view, which substantially reduces the demand for storage and memory capacity during further processing (average reduction of 16,170 processed fluorescently labelled body images: 13.4-fold).

### Automatic quality control of image acquisition and processing improves measurement precision

Outliers arising in image quality or sporadic failures in automated analysis can distort quantitative trajectories in longitudinal imaging experiments. To address this, ALIGN implements an automated quality control (QC) module that identifies and removes technical outliers prior to downstream analysis (Fig. 4).

**Figure 4:**
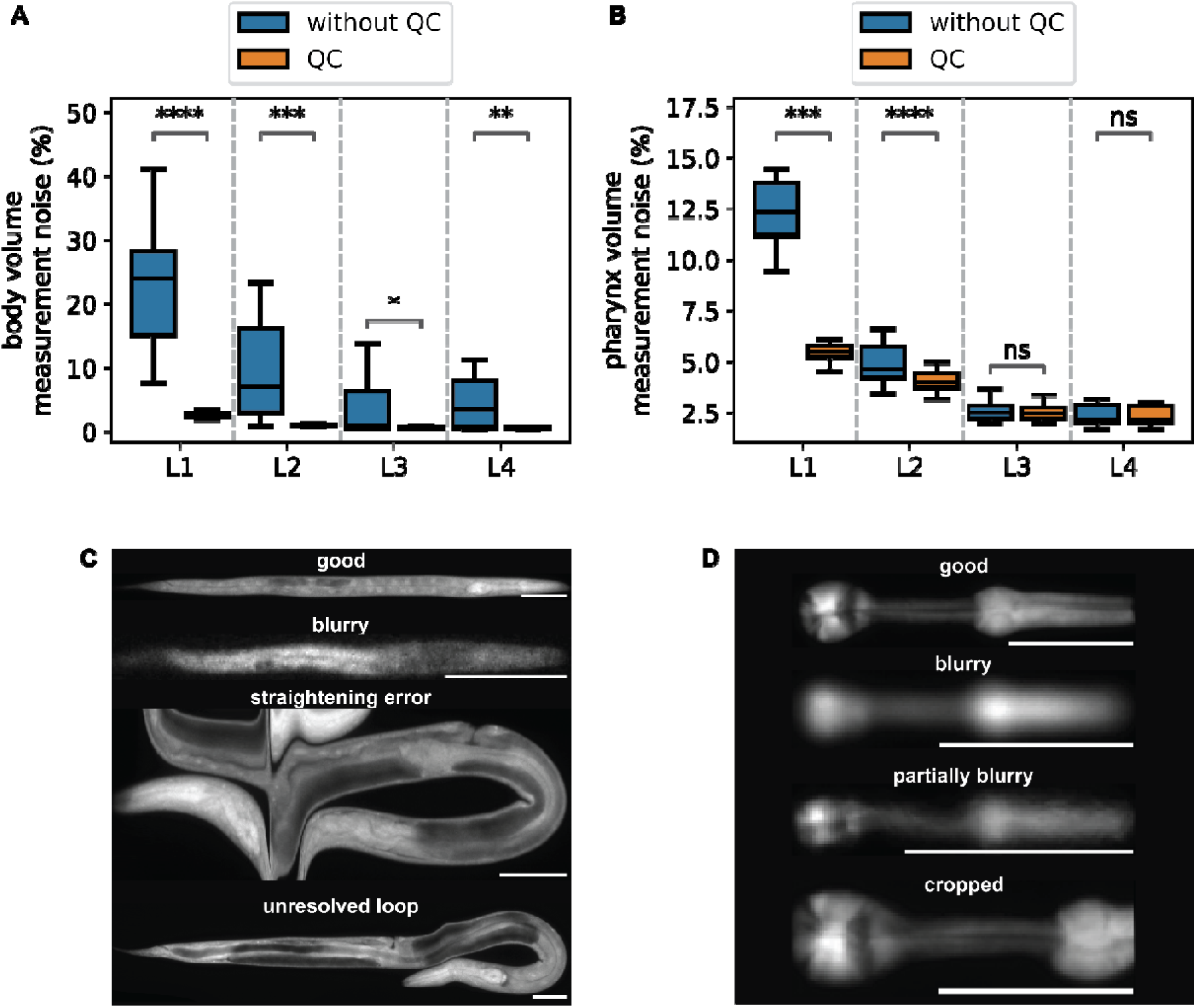
Automated quality control improves measurement precision. **(A)** Boxplot of the coefficient of variation among repeated measurements for the whole-body volume with and without automated quality control. The time series of individuals of each larval stage were rescaled to a constant length (and matching moult entry and exit points) and averaged over each larval stage. (n=30 individuals, 98 time points per individual, 20 measurements per individual and time point, from a single experiment). (Wilcoxon signed-rank test, two-sided, Holm-Bonferroni correction) **(B)** As (A), but for pharynx size (Wilcoxon signed-rank test, two-sided, Holm-Bonferroni correction) **(C)** Examples of accepted and rejected images of straightened worms expressing a somatic mScarlet marker (*eft-3p*::mScarlet). **(D)** Examples of accepted and rejected images of straightened pharynxes (*myo-2p*::GFP). Scale bar: 50 µm. *P<0.05; **P<0.01; ***P<0.001; ****P<0.0001; ns, non-significant. Exact p-values provided in Supplemental Table 2. Boxplots: centre line, median; box, interquartile range (IQR); whiskers, 1.5×IQR.

QC classification is based on features extracted from straightened images and corresponding segmentation masks using region-based descriptors, followed by classification with gradient-boosted decision trees (XGBoost) (Chen and Guestrin, 2016). The default model detects common failure modes observed in the datasets of longitudinal *C. elegans* imaging, including poor focus, segmentation errors affecting straightening, and unresolved body postures such as loops or omega turns (Fig. 4C, 4D). Importantly, the QC framework can be readily adapted to new datasets using a dedicated napari-based annotation interface, enabling users to train dataset-specific classifiers (Supplemental Movie 3).

To quantify the impact of QC on measurement precision, we generated a reference dataset consisting of images from repeated acquisitions at each time point under identical conditions. We used a strain ubiquitously expressing a red fluorescent protein (eft-3p::mScarlet) and a green fluorescent protein specifically expressed in the pharyngeal muscle (myo-2p::GFP) (Stojanovski et al., 2023). Measurement noise in body and pharynx size was estimated as the coefficient of variation (CV) across repeated measurements spaced by ∼10 seconds. For this analysis, we assume that biological change over the acquisition interval is negligible, such that any remaining variation between time points represents measurement noise.

Application of QC reduced measurement variability for both body and pharyngeal size (Fig. 4A-4B), with the strongest improvement observed in early larval stages, where the signal was weakest, and where image processing was most sensitive to out-of-focus blur. This increase in precision was achieved without substantially reducing the measurement time resolution (removed points: 11.3% for body measurements, 20% for pharynx measurements), indicating efficient discrimination between technical outliers and valid measurements. Notably, the tradeoff between retaining the maximal number of time points and reaching optimal precision can be tuned by training the QC model to what the user considers acceptable image quality for a given experimental design.

These results demonstrate that automated QC is instrumental for robust quantitative analysis of longitudinal imaging data and can be flexibly adapted to different imaging conditions and target structures.

### Automated detection of larval moults from growth trajectories

During post-embryonic development, *C. elegans* progresses through four larval stages separated by lethargus, a phase of growth arrest, followed by cuticular ecdysis during larval moults (Singh and Sulston, 1978). These dynamics are reflected in body size trajectories as characteristic growth plateaus followed by renewed growth, providing a basis for automated detection of larval stage transitions (Olmedo et al., 2015; Singh and Sulston, 1978). However, precise identification of moult timing is challenging, as transition points are sensitive to noise in the growth trajectory, and irregular growth patterns can lead to false positives.

To robustly identify moult events, we trained a one-dimensional U-Net model to detect larval stage transitions directly from body size trajectories using over ten thousand manually annotated ecdysis events as ground truth. This approach enables automated prediction of moult timing with high accuracy (average error = 0.8 time points ≈ 8 minutes) and a low failure rate (misdetection rate = 2.5%; over-detection rate = 3.7%) (Fig. S2).

For validation and refinement of these automated moult assignments, ALIGN integrates an interactive graphical interface that overlays predicted moult events onto growth trajectories and links them to the corresponding image data. This interface allows users to efficiently inspect predictions, correct misassigned events, and annotate additional features within a unified environment. The interface provides structured access to large time series datasets at defined developmental stages, enabling rapid exploration of longitudinal phenotypes (Supplemental Movie 4).

### Example application: ALIGN quantifies growth and feeding dynamics in *eat-2* mutants

To demonstrate the utility of ALIGN for integrative longitudinal analysis, we quantified feeding behaviour, growth, and body size in *eat-2(ad1113)* mutants. In this mutant, pharyngeal pumping is perturbed due to a mutation in the acetylcholine receptor involved in neuromuscular signal transduction (Avery, 1993; Raizen et al., 1995). Mutations in *eat-2* have been reported to reduce pumping rates and growth (Avery, 1993). However, these measurements have largely relied on manual scoring (Raizen et al., 2012), been applied to immobilised worms (Scholz et al., 2016), or on single time-point measurements of adults (Bonnard et al., 2022), and have not tracked individuals longitudinally during their development.

Using agarose microchambers, we imaged individual wild-type and *eat-2* mutant animals expressing a ubiquitous mScarlet marker and a pharyngeal GFP reporter from hatching to the fourth larval moult. Time-lapse imaging was performed at 20-minute intervals. In addition, we acquired high-frame-rate recordings (for 8 s at 25 frames per second) hourly to quantify pharyngeal pumping dynamics.

Consistent with previous reports, *eat-2* mutants exhibited reduced pharyngeal pumping rates, decreased body size, and slower growth across development (Fig. 5A, 5C). In addition to these population-level effects, ALIGN enabled analysis of individual trajectories and revealed increased variability in feeding behaviour. Specifically, pumping in *eat-2* mutants was less regular, with more frequent non-oscillatory pauses compared to wild-type animals (Fig. 5E-G).

**Figure 5:**
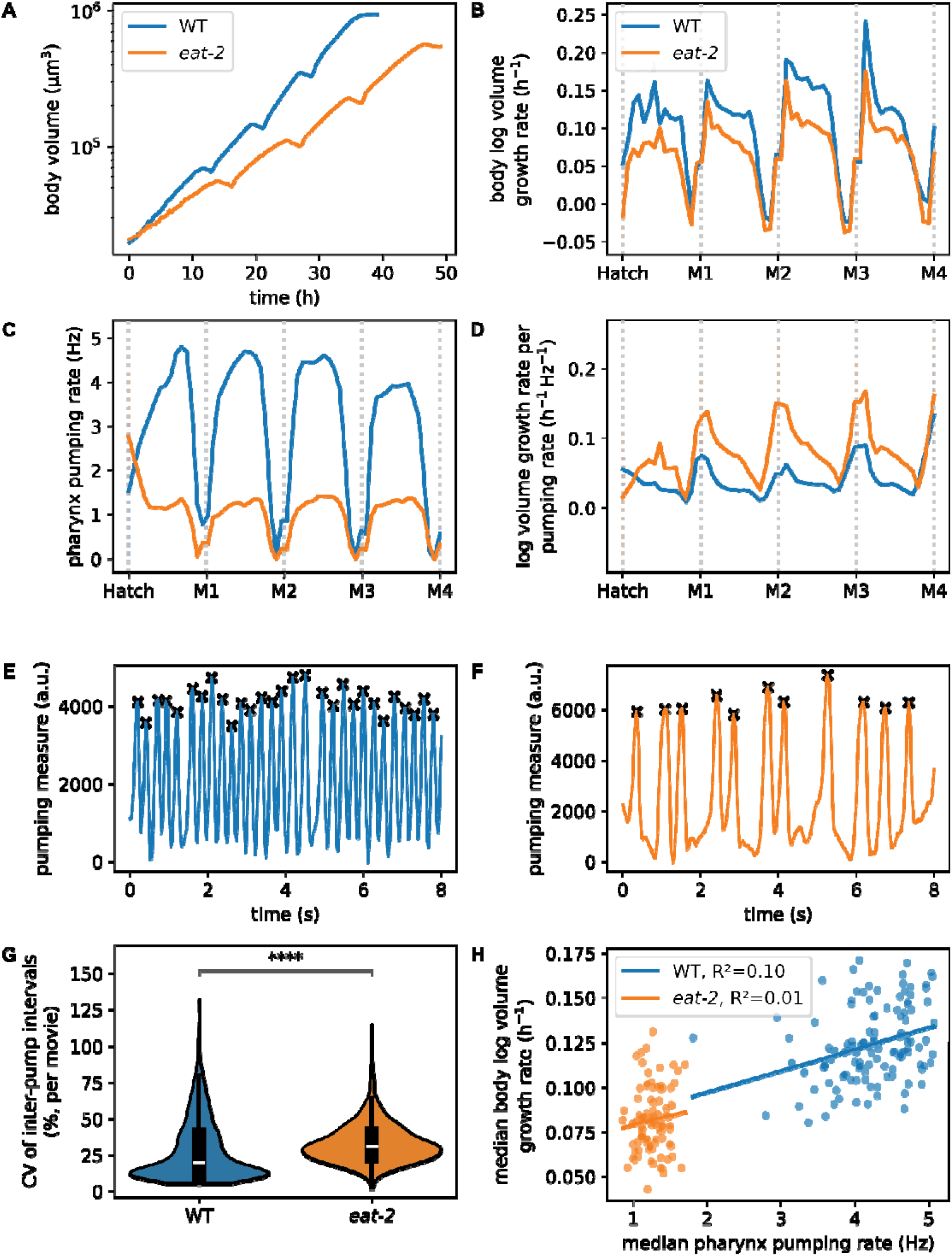
ALIGN reveals individual longitudinal information on pharyngeal pumping in wild type and *eat-2* mutants. **(A)** Body volume as a function of time averaged from n=37 (WT) and n=25 (*eat-2* mutant) animals. For averaging, the time series of individuals of each larval stage were rescaled to a constant length (matching moult entry and exit points). After averaging, the series was scaled back to the mean larval stage duration of the corresponding condition. (Shaded area = 95% Confidence Interval (CI)). **(B)** Exponential body volume growth rate (dlog(V)/dt) as a function of developmental progression from n=37 (WT) and n=25 (*eat-2* mutant). For each larval stage, growth rate series of individuals were rescaled to a constant length. The four larval stages were then concatenated with each stage scaled to be 25% of the horizontal axis (i.e., M1-M4 occur at 25%, 50%, 75%, and 100%). (Shaded area = 95% CI). **(C)** Pharyngeal pumping rate as a function of developmental progression. (Shaded area = 95% CI). **(D)** Ratio of exponential body volume growth rate and pharyngeal pumping rate as a function of developmental progression. (Shaded area = 95% CI). Movies where no pumps were detected were removed from the calculation. **(E)** Representative pumping index obtained from the analysis of 25 FPS movies of a wild-type animal. Crosses indicate detected pumping events. **(F)** As (E), but for an *eat-2* mutant animal. **(G)** Violin and overlaid boxplot of the coefficient of variation of inter-pump interval. CV was determined for every movie individually. Movies with fewer than 5 detected pumping events were removed from the analysis to ensure meaningful CV computation (n=1870 (WT), n=1413 (*eat-2*), Mann-Whitney-Wilcoxon test, two-sided). **(H)** Scatterplots of the median pumping rate against the median log body volume growth rate and the linear regression fit of each population. For each individual, each series was rescaled as in (B). 20% of the series of each larval stage was trimmed at the beginning and at the end to avoid artefacts caused by rapid volume expansion during moulting. Each point represents a unique worm-stage pair. (Shaded area = 95% CI of the linear regression, WT: β = 0.0125, ε = 0.0715, R² = 0.10, p = 1.0958e-04, n = 118; *eat-2*: β = 0.01, ε = 0.0676, R² = 0.0126, p = 2.9479e-01, n = 89). ****P<0.0001. Exact p-values provided in Supplemental Table 2. Boxplots: centre line, median; box, interquartile range (IQR); whiskers, 1.5×IQR.

Notably, although pumping rates were reduced by approximately four-fold in *eat-2* mutants (Fig. 5C), growth rate decreased by less than two-fold (Fig. 5B, 5D). Similarly, pumping frequencies correlated only weakly with the growth rates of individual animals (R^2^ =0.10 for wild type and R^2^ =0.01 for *eat-2*) (Fig. 5H). Together, these data suggest that organi smal growth is not solely limited by pumping frequency and instead depends on additional factors such as the efficiency of nutrient utilisation.

Together, these results demonstrate that ALIGN enables automated, longitudinal, and multi-modal quantification of behaviour and growth at single-animal resolution. This approach was validated by confirming the reduced pharyngeal pumping rate of *eat-2* mutants. Going beyond validation, ALIGN was used to additionally reveal previously inaccessible relationships between feeding dynamics and growth. The methodology thereby provides a framework for studying longitudinal behavioural and growth heterogeneity throughout development at high throughput and precisi on.

## Discussion

We present ALIGN, a modular framework for reproducible and scalable analysis of longitudinal microscopy data that integrates conventional image processing, deep learning-based segmentation, and interactive curation within a unified workflow. By combining automated analysis with visualisation and manual refinement, ALIGN enables robust extraction of quantitative features from complex time-lapse datasets. Additionally, the establishment of an animal-centred coordinate system allows consistent measurement of morphology and dynamics in motile specimens, including reconstruction of three-dimensional structures from misaligned image stacks.

Using this framework, we quantified feeding behaviour, growth, and size in *C. elegans* eat-2 mutants across development, demonstrating that ALIGN facilitates simultaneous, longitudinal analysis of multiple phenotypic dimensions at single-animal resolution. Beyond reproducing previously reported phenotypes, this approach revealed increased variability in feeding dynamics and a non-proportional relationship between pumping rate and growth, illustrating how integrated analysis can uncover relationships that are difficult to access with conventional methods.

A central feature of ALIGN is its accessibility to non-specialist users. Workflow definition through a simple configuration file allows for reproducible execution without custom programming, while the modular block-based archi tecture ensures transparency and extendibility. At the same time, the framework supports large-scale data processing through integration with distributed computing environments, allowing efficient analysis of datasets comprising hundreds of thousands of images. Standardised data structures further facilitate downstream analysis and integration with external tools.

A prominent challenge of segmentation and image processing pipelines lies in their sensitivity to imaging conditions. ALIGN addresses this challenge by providing coding-free, interactive interfaces for creating training data and for retraining segmentation networks. Nevertheless, usage of ALIGN on other types of data, or with different imaging conditions might requi re minor adjustment to the code, such as algorithm customisation. The current implementation of the straightening algorithm is suitable for elongated or tubular structures in general, but we have only thoroughly tested i t for *C. elegans* body and pharynx. Adaptation to more complex morphologies may require refinement of straightening and alignment strategies.

The modular design of ALIGN facilitates extension both to further applications for *C. elegans* and to other study systems. Specifically, the framework can be adapted to other longitudinal imaging systems, and its integration of automated analysis with interactive curation provides a general strategy for combining machine learning with human validation. In particular, the reconstruction of 3D information of moving specimens by realignment of individual planes will likely find application in other setups and species. To date, the standard approach for imaging moving specimens is physical immobilisation (Berger et al., 2021; Keil et al., 2017), which often involves complex experimental designs that do not scale well and face the risk of perturbing specimen physiology. Our computational approach offers an effective and transferable solution to this common problem in live imaging of moving organisms. The only constraints of the method are that the specimen must not deform during acquisition, except for tubular bending, and that movement in the axial direction must be negligible.

The bottleneck in longitudinal imaging has shifted from acquisition to analysis and data management. ALIGN covers the full path from raw images to curated trajectories in one reproducible workflow. It exposes every step of this path to the user, including retraining of the segmentation and quality-control models, allowing flexible adjustment to the specifics of a given experimental setting. The workflow is intended to make single-animal developmental trajectories a routine measurement rather than a specialist undertaking, including in laboratories without dedicated computational experti se. We expect its principal value to emerge where the scale of imaging outgrows what can be curated and tuned by hand, in particular in longitudinal experiments following large numbers of individuals. In this regime, variation between individuals and over time, long treated as a nuisance parameter in developmental biology, becomes an accessible object of study.

## Materials and methods

### Software language and packages

ALIGN consists of a Python package available under https://doi.org/10.5281/zenodo.22935167 and distributed using pip, and a pipelining tool: https://doi.org/10.5281/zenodo.22935125. Both are available as free and open-source software under the BSD-3-Clause licence. During installation, ALIGN creates a dedicated micromamba (QuantStack and mamba contributors, 2026) environment. Validated package and dependency versions are maintained using lock files created with conda-lock (conda-lock contributors, 2026). See https://spsalmon.github.io/align_pipeline/ for installation and usage guides. Interactive tools include a time series annotator, developed using Shiny for Python (The Shiny development team, 2026) and a napari (napari contributors, 2019) plugin (available at https://doi.org/10.5281/zenodo.22935002). A small sample dataset is available here: https://doi.org/10.5281/zenodo.22939939. Pre-trained models for segmentation, quality control and moult detection are available here: https://doi.org/10.5281/zenodo.23038693.

### Use of AI tools

Large Language Models (LLMs) such as Claude and ChatGPT were used for the preparation of this manuscript. Their use included: generation of parts of the code, code refactoring, rewriting of parts of this manuscript (typo and error detection and correction, reformulation of sentences, etc.). LLMs were always used under careful human supervision.

### Strains

*C. elegans* strains shown in this study were created by genetic crosses from the following previously published and extensively outcrossed alleles: bqSi577 [*myo-2p::gfp*] (GFP-tagged pharynx) (Muñoz-Jiménez et al., 2017), wbmIs88 [*eft-3p::3xflag::dpy-10 crRNA::sl2::wrmScarlet::unc-54* 3ü UTR] (somatic mScarlet) (Silva-García et al., 2019), sam124 [*glh-1::T2A::sGFP2*] (GFP-tagged germline) (Goudeau et al., 2021), [Y75B12B.1p::nls::ce-gfp::nls::tbb-2 3ü UTR] (GFP-tagged germline nuclei) (Aljohani et al., 2023), bqSi225 [pBN34(unc-119(+) Pemr-1::emr-1::mcherry)] (mCherry-tagged nuclear envelope) (Morales-Martínez et al., 2015), and *eat-2(ad1113)*.

List of strains used:

• wBT125: *bqSi577 [myo-2p::GFP + unc-119(+)] IV; wbmIs88 [eft-3p::3xFLAG::dpy-10 crRNA::SL2::wrmScarlet::unc-54 3Ö UTR] V:8645000 (1. 52); Fig. 1; Fig. 2B, 2C, 2D; Fig. 3A, 3B, 3C; Fig. 4; Fig. 5*.

• wBT318: *rpl-34(syb1880) [rpl-34:RPL-34:mCherry] IV:5. 83; glh-1(sam124[glh-1::T2A::sGFP2]) I:1. 41; Fig. 2E*.

• wBT396: *bqSi225[pBN34(unc-119(+) Pemr-1::emr-1::mcherry)] IV; Fig. 2F*.

• wBT674: *wbmIs88[eft-3p::3xFLAG::dpy-10 crRNA::SL2::wrmScarlet::unc-54 3Ö UTR] V:8645000 (1. 52); kstSi148[cbr-unc-119(p1+p2, loxP)(+), Y75B12B. 1p::nls::ce-gfp::nls::tbb-2 3Ö UTR] II; unc-119(ed3) III; Fig. 3D*.

• wBT615: *eat-2(ad1113) II; bqSi577 [myo-2p::GFP + unc-119(+)] IV; wbmIs88 [eft-3p::3xFLAG::dpy-10 crRNA::SL2::wrmScarlet::unc-54 3Ö UTR] V:8645000; Fig. 5*.

### Agarose-based cultivation chamber preparation

Agarose-based chambers were assembled as previously described (Turek et al., 2015) with minor adaptations (Stojanovski et al., 2022). In brief, a polydimethylsiloxane (PDMS) master mold was used to pattern wells in an S-basal 4.5% agarose gel, which were filled with OP50-1 bacteria as food. After filling wells with embryos, chambers were sealed by inversion on a gas-permeable high optical quality polymer mounted on a 3.5 cm dish (ibidi), surrounded by 3% low-melting temperature agarose and covered with PDMS. Chamber dimensions were 600×600×20 µm (Fig. 1; Fig. 2B-E; Fig. 3A-C; Fig. 4; Fig. 5), 200×200×10 µm (Fig. 2F), and 350×350×10 µm (Fig. 3D, 3E, 3G, 3H) matching the field of view of the objective used. Dimensions are given as width x height x depth.

### Imaging

Experiments of Figure 1F and experiments shown in Supplemental Fig. 1 were acquired using a SQUID microscope (Cephla) using a 10x Olympus objective (NA = 0.4). Images shown in Figures 2F and 3D were acquired using a Nikon Ti2 microscope equipped with a W1 Yokogawa spinning disc scan head, a Lumencor ZIVA light engine, and a dual Photometrics Kinetix22 CMOS camera at 60x magnification (NA=1.4, oil immersion). The experiment featured in Figure 5 was acquired using epifluorescence mode on a Nikon Ti2 microscope (Photometrics Prime BSI CMOS camera, 10x Nikon air objective (NA=0.45), Lumencor SpectraX light source). All other images shown in the main text were acquired using a Nikon Ti2 epifluorescence microscope (Hamamatsu ORCA Flash 4 sCMOS camera, 10x Nikon air objective (NA=0.45)).

### Pipeline inputs

Input files were provided to the ALIGN pipeline as individual ome.tiff files with filenames containing information on time and a position identifier. The exact input pattern can be defined in the configuration file (see documentation https://spsalmon.github.io/align_pipeline/). Each individual file was a multi-page TIFF containing all channels, Z-planes and time frames (in case of pharyngeal pumping videos).

### Deep learning segmentation

Deep learning models were implemented using PyTorch (Paszke et al., 2019). For network training, a dataset of images was selected that spans all developmental stages, experimental conditions, and imaging modalities by a Python script included in the ALIGN package, ensuring an appropriate balance of these different image classes. Ground truth segmentation masks were subsequently annotated and curated using the microSAM napari plugin (Archit et al., 2025).

Model architecture was defined through the segmentation_models.pytorch package (Iakubovskii, 2019). U-Net++(Zhou et al., 2018) architecture with an efficientnet-b5 encoder (Tan and Le, 2019) trained on ImageNet (Deng et al., 2009) was used for all networks shown, which superseded other architecture designs tested. The ADAM (Kingma and Ba, 2017) optimiser was used with a learning rate of 10⁻⁴. Focal Tversky loss (Abraham and Khan, 2019) was used to adjust the weight of otherwise underrepresented foreground pixels.

Images were preprocessed by percentile-based normalisation per image and channel using default values spanning between the 1^st^ and 99^th^ percentiles. These normalisation parameters are included as hyperparameters of the model and are automatically reused during prediction.

All training parameters, such as architecture, normalisation, loss function, learning rate, batch size, and train-test split ratio are specified in a YAML configuration file. This file and dataframes of the training, validation, and test sets are automatically copied into the file path of the model during training. Size of training, test, and validation sets for pre-trained models developed here are li sted in Supplemental Table 1.

### Computational straightening

The straightening algorithm comprised the following steps, using an image of the object to be straightened and the corresponding segmentation mask as an input, building on previous approaches (Peng et al., 2008).

A medial axis transform (MAT) was used to extract the morphological skeleton. Branches of the skeleton were removed by selecting the longest shortest path and trimming the ends by the mask’s radius (defined as 50% of the maximum value of the distance transform of the mask) to remove frequently occurring wrongly skeletonised regions near the endpoints. To fill in the trimmed end sections, two perpendicular rays were projected to either side of the midline and their intersection with the mask border was determined. The midpoint between these intersecti ons along the mask border was added to the midline. Cubic splines were used to smooth and interpolate the midline, as well as fill the gap between skeleton and endpoints. Interpolation was extended by 1% beyond the endpoints. Finally, the smoothed spline was used to compute the 2D coordinate map orthogonal to and along the midline. For presentation and feature extraction, images and masks were warped to those midline-centred coordinates.

When applied to 3D stacks or videos, splines and straightening were computed for each plane. Straightened planes were then iteratively aligned using Phase Cross Correlation (Guizar-Sicairos et al., 2008). The parameterisation of the splines was then offset, and the set of final splines was fit based on the updated parameterisation.

### Feature extraction

Length was computed from the straightened image as the number of non-zero columns along the x axis of the straightened mask. Volume was inferred from 2D images, assuming rotational symmetry along the mask’ s x axis. Area was calculated from the non-straightened mask as the sum of all non-zero pixels. All measurements were converted to physical units using the pixel size defined by the user.

### Automated image quality control and classification

To classify images as worms, eggs or errors, two consecutive binary XGBoost classifiers were used. First, images of eggs were detected based on the output of full body segmentation. Second, remaining images were split into correctly segmented and straightened worms and errors. For quality control of organ segmentation, the same procedure was applied after removing images classified as eggs (based on the full body segmentation) to fl ag poor quality images.

Classifiers were trained using straightened images and masks. Raw images were normalised as described under segmentation. All features available in the regionprops function of scikit-image (van der Walt et al., 2014) as well as the variance of the image divided by its mean were used for model training. Hyperparameters of the models (such as maximum depth or learning rate) were optimised using Bayesian Optimisation (as implemented in the BayesianOptimization package (Nogueira, 2014)). Training datasets were created as described under segmentation and manually annotated using a napari plugin provided in the package.

### Automated moult detection

For creating a ground truth dataset, moults were manually annotated in volume trajectories for 11,245 individuals. For each individual, every moult was represented as a separate output channel. A one-dimensional Gaussian centred on the frame at which the moult occurred was generated over the full trajectory, with a fixed standard deviation of 2 frames. Moults that were absent for a given individual were assigned an all-zero target channel. A 1D U-Net was trained to reproduce these Gaussians from raw trajectory data. For each individual, moult existence was predicted independently of moult timing by an auxiliary presence head. The final decoder feature map was reduced to a single feature vector per individual by averaging over time. Padded frames were excluded from this average. This vector was passed through a two-layer perceptron to produce one logit per moult, representing the probability that the corresponding moult was present in the trajectory. Learning rate was 10⁻⁴. For the loss function, a peak-weighted mean squared error loss function was used for the heatmap regression and was combined with a Binary Cross Entropy Loss for event presence detection, with the formula:

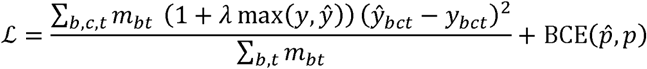

where *b, c, t* each represent the batch, channel and time dimension and m is a mask of all non-padded values. The weight coefficient *λ* was set to 3 to emphasise accurate prediction around the events as the heatmaps are dominated by near-zero values. Individual moults were called at peaks of the predicted Gaussians if the moult was predicted to be present in the series.

### Quantification of pumping rate

Movies of worms expressing the pharyngeal marker *myo-2p::GFP* were acquired at 25 frames per second. For each movie, all frames were segmented, straightened, and aligned. Pixel intensities were normali sed to span between 0 and 1. The position of the terminal bulb (where the pharynx is the widest and pharyngeal muscle movement is strongest) was identified by the peak of the intensities summed along the vertical axis (i.e., orthogonal to the body axis) furthest away from the middle of the straightened image. The pumping index was calculated as the negative standard deviation of the normali sed intensity of pixels located at the terminal bulb, following previous approaches (Bonnard et al., 2022).

The pumping index time series was detrended by subtracting a Savitzky-Golay-filtered version of the signal (window length = 101 frames, polynomial order = 5) to remove baseline drift while preserving the higher-frequency pumping events. The detrended signal was then smoothed using a Whittaker smoother (*λ* = 0.25, second-order differences) to reduce residual noise and shifted to non-negative values by adding the absolute value of its minimum. Pumping events were detected as peaks in this processed signal using scipy.signal.find_peaks, with constraints on minimum inter-peak distance, peak width, peak height, and a prominence threshold set to a fixed multiple of the signal’s standard deviation. Peak selection criteria were manually optimised for a given dataset. The pumping rate was computed as the number of detected peaks divided by the total recording duration (in seconds), yielding pumps per second.

## Acknowledgements

We thank Peter Lenart, Ioana Gheorghe, and Klement Stojanovski for providing images, and Guillaume Witz, Maciej Dobrzinsky, and Roman Schwob for critical reading and helpful feedback on the manuscript. We acknowledge support by the Microscopy Imaging Center at the University of Bern. Some strains were provided by the CGC, which is funded by NIH Office of Research Infrastructure Programs (P40 OD010440). This research was funded in part by the Swiss National Science Foundation (SNSF) [grants PCEFP3_181204, 310030_219822, 320030L-227534]. For the purpose of open access, a CC BY public copyright licence is applied to any Author Accepted Manuscript (AAM) version arising from this submission.

## Data and resource availability

All code developed in this project is available in public repositories listed under “Software language and packages” in material s and methods.

