## Supplemental Information for "ALIGN: A modular framework for quantitative analysis of longitudinal development in *Caenorhabditis elegans*"

- **Supplemental Figure 1**
- **Supplemental Figure 2**
- **Supplemental Table 1**
- **Supplemental Table 2**
- **Supplemental Movie 1**
- **Supplemental Movie 2**
- **Supplemental Movie 3**
- **Supplemental Movie 4**

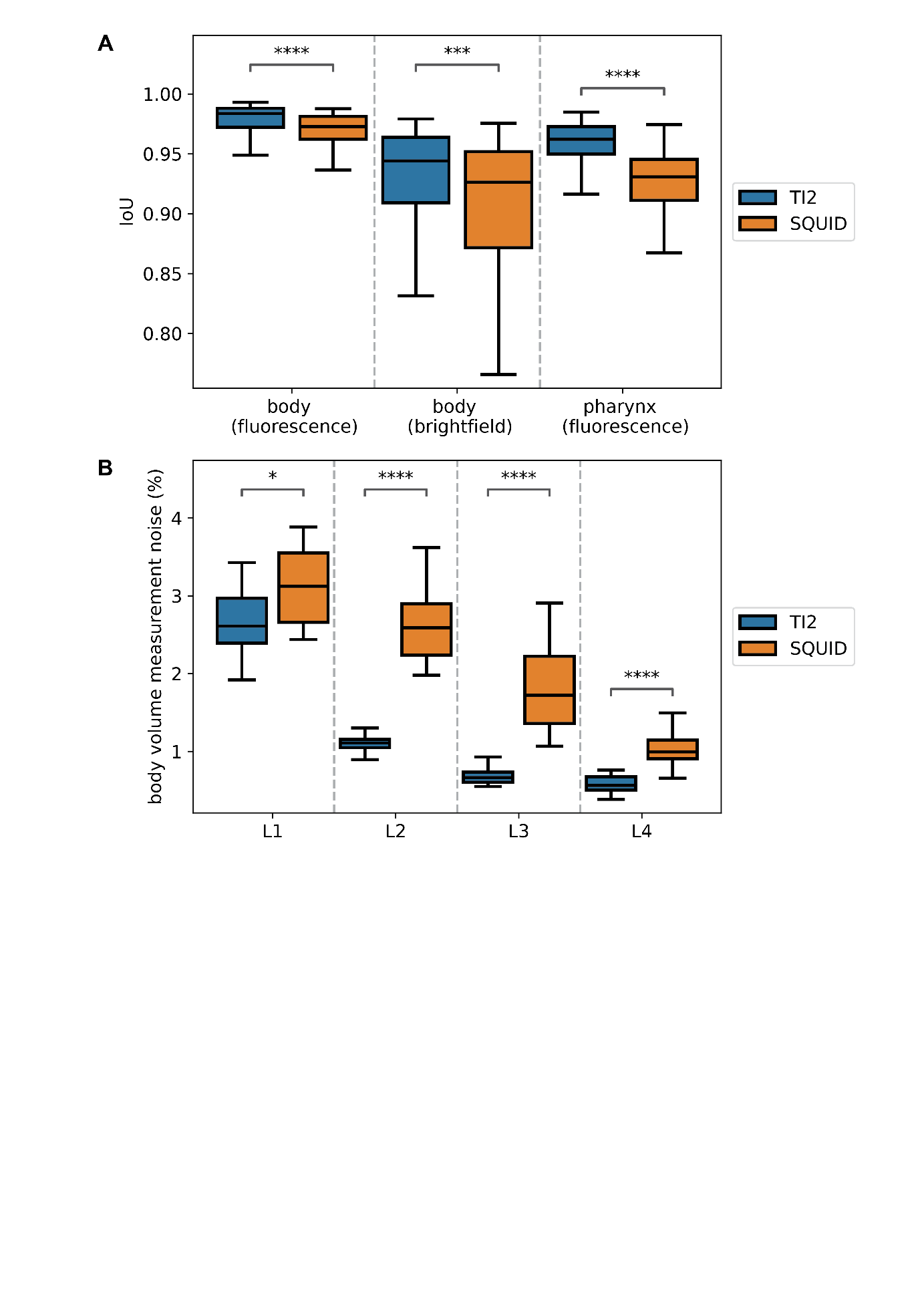

**Supplemental Figure 1: Comparison of ALIGN’s performance between a TI2 widefield microscope and a SQUID microscope.
A.** Boxplot of segmentation accuracy for images acquired on a TI2 microscope and on a SQUID microscope. (TI2: body=117 images, body (brightfield)=204 images, pharynx=185 images; SQUID: body=37 images, body (brightfield)=91 images, pharynx=91 images). (Mann-Whitney-Wilcoxon, two-sided, Holm-Bonferroni correction). **B.** Boxplot of the coefficient of variation among repeated measurements for the whole-body volume, from an experiment acquired on a TI2 microscope and one acquired on a SQUID microscope. (TI2: n=30 individuals, 98 time points per individual, 20 measurements per individual and time point. SQUID, experiment 1: n=24 individuals, 336 time points per individual, 10 measurements per individual and time point. SQUID, experiment 2: n=21 individuals, 140 time points per individual, 10 measurements per individual and time point). (Mann-Whitney-Wilcoxon, two-sided, Holm-Bonferroni correction). *P<0.05; ***P<0.001; ****P<0.0001. Exact p-values provided in Supplemental Table 2. Boxplots: centre line, median; box, interquartile range (IQR); whiskers, 1.5×IQR.

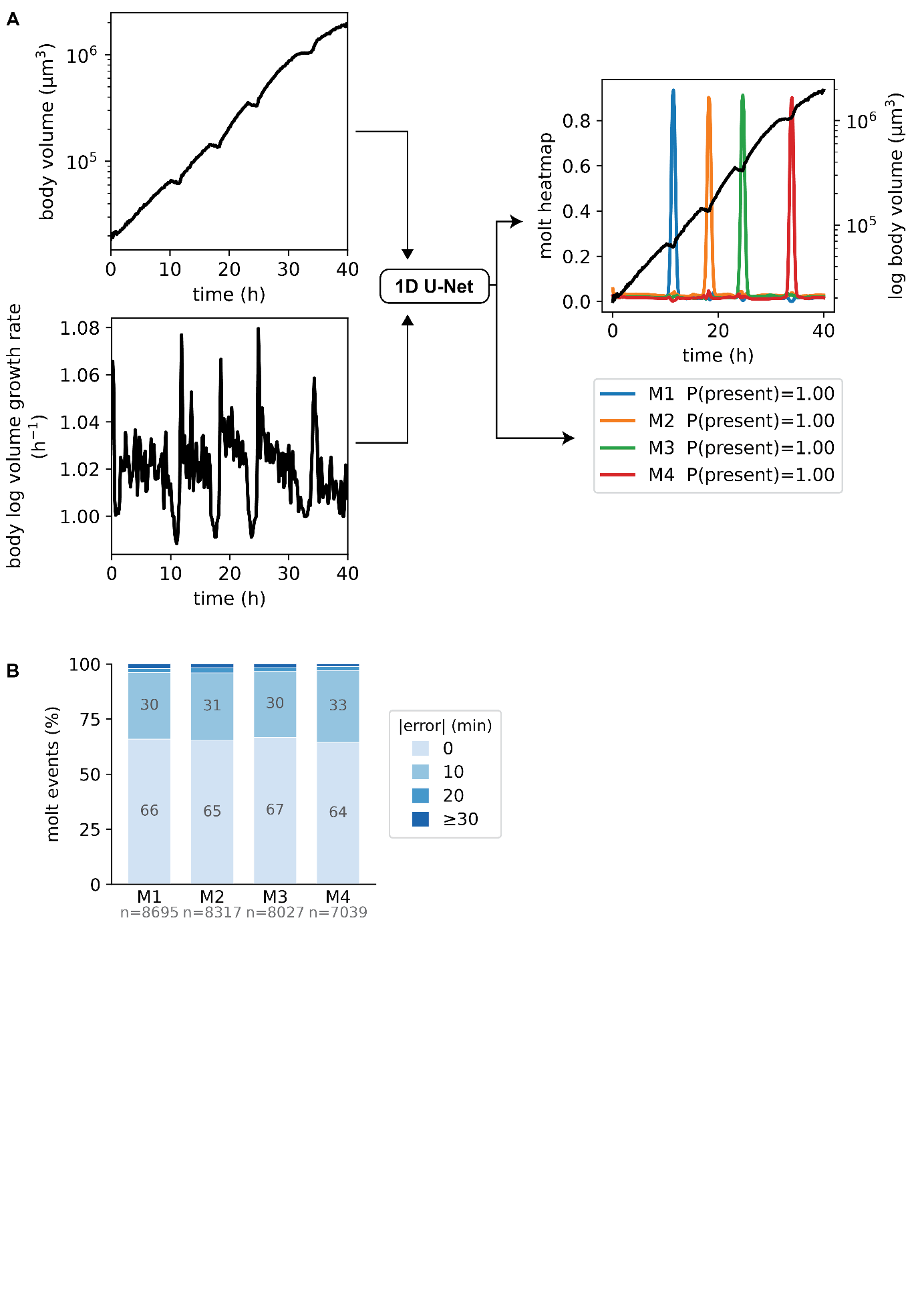

**Supplemental Figure 2: Automated moult detection using a 1D U-Net.
A.** For a given individual, using a series of its body volume and body volume growth rate over time, we predict the position of the moults using a 1D U-Net. A prediction head (typical U-Net output) predicts a 1D heatmap for each moult while a presence head (a two-layer MLP, Linear-ReLU-Linear) predicts if each moult is present in the series or not. **B.** Proportions of the prediction errors for each moult event in the test dataset.

|  | Training set size | Validation set size | Testing set  size |
| --- | --- | --- | --- |
| Whole body (brightfield) | 2,059 | 588 | 295 |
| Whole body (fluorescence) | 1,397 | 399 | 201 |
| Pharynx (fluorescence) | 1,924 | 550 | 276 |
| Germline (fluorescence) | 1,337 | 382 | 192 |

**Supplemental Table 1: Number of images for each set used for training the provided segmentation models.**

| Figure | Panel | Comparison | Test | p-value |
| --- | --- | --- | --- | --- |
| 2 | C | DL (fluo) vs edge (fluo) | Wilcoxon signed-rank test, two-sided | 2.207e-34 |
| 2 | C | DL (BF) vs edge (fluo) | Mann-Whitney-Wilcoxon, two-sided | 2.642e-02 |
| 2 | D | DL vs edge | Wilcoxon signed-rank test, two-sided | 2.102e-35 |
| 2 | E | DL vs edge | Wilcoxon signed-rank test, two-sided | 6.747e-31 |
| 3 | F | Raw vs straightened | Wilcoxon signed-rank test, two-sided | 1.869e-19 |
| 4 | A | No QC vs QC (L1) | Wilcoxon signed-rank test, two-sided, Holm-Bonferroni correction | 6.104e-05 |
| 4 | A | No QC vs QC (L2) | Wilcoxon signed-rank test, two-sided, Holm-Bonferroni correction | 4.378e-04 |
| 4 | A | No QC vs QC (L3) | Wilcoxon signed-rank test, two-sided, Holm-Bonferroni correction | 1.252e-02 |
| 4 | A | No QC vs QC (L4) | Wilcoxon signed-rank test, two-sided, Holm-Bonferroni correction | 3.346e-03 |
| 4 | B | No QC vs QC (L1) | Wilcoxon signed-rank test, two-sided, Holm-Bonferroni correction | 1.221e-04 |
| 4 | B | No QC vs QC (L2) | Wilcoxon signed-rank test, two-sided, Holm-Bonferroni correction | 3.815e-06 |
| 4 | B | No QC vs QC (L3) | Wilcoxon signed-rank test, two-sided, Holm-Bonferroni correction | 4.311e-02 (ns after correction) |
| 4 | B | No QC vs QC (L4) | Wilcoxon signed-rank test, two-sided, Holm-Bonferroni correction | 2.850e-01 |
| 5 | G | WT vs *eat-2* | Mann-Whitney-Wilcoxon, two-sided | 8.081e-51 |
| S1 | A | TI2 vs SQUID (body, fluo) | Mann-Whitney-Wilcoxon, two-sided, Holm-Bonferroni correction | 4.737e-05 |
| S1 | A | TI2 vs SQUID (body, brightfield) | Mann-Whitney-Wilcoxon, two-sided, Holm-Bonferroni correction | 1.619e-04 |
| S1 | A | TI2 vs SQUID (pharynx, fluo) | Mann-Whitney-Wilcoxon, two-sided, Holm-Bonferroni correction | 7.739e-26 |
| S1 | B | TI2 vs SQUID (L1) | Mann-Whitney-Wilcoxon, two-sided, Holm-Bonferroni correction | 1.678e-02 |
| S1 | B | TI2 vs SQUID (L2) | Mann-Whitney-Wilcoxon, two-sided, Holm-Bonferroni correction | 1.012e-07 |
| S1 | B | TI2 vs SQUID (L3) | Mann-Whitney-Wilcoxon, two-sided, Holm-Bonferroni correction | 1.094e-07 |
| S1 | B | TI2 vs SQUID (L4) | Mann-Whitney-Wilcoxon, two-sided, Holm-Bonferroni correction | 3.108e-06 |

**Supplemental Table 2: Statistical tests and p-values of the different comparisons.**

**Supplemental Movie 1:** Montage of the segmentation masks of 100 WT C. elegans during their entire larval development. Worms expressing a bright somatic red fluorophore (eft-3p::mScarlet) were imaged every 10 minutes over two days and segmented using a deep learning model in ALIGN.

**Supplemental Movie 2:** Montage of images of 25 WT *C. elegans* expressing a bright somatic red fluorophore (*eft-3p*::mScarlet) during their entire larval development, straightened using ALIGN (time resolution: 10 minutes).

**Supplemental Movie 3:** Usage example of ALIGN’s napari plugin for data annotation, applied to a classification task.

**Supplemental Movie 4:** Usage example of ALIGN’s GUI linking time series and source images, for exploring and annotating time series (moults, death, etc.).
